# Inactive USP14 and inactive UCHL5 cause accumulation of distinct ubiquitinated proteins in mammalian cells

**DOI:** 10.1101/479758

**Authors:** Jayashree Chadchankar, Victoria Korboukh, Peter Doig, Steve J. Jacobsen, Nicholas J. Brandon, Stephen J. Moss, Qi Wang

## Abstract

USP14 is a cysteine-protease deubiquitinase associated with the proteasome and plays important catalytic and allosteric roles in proteasomal degradation. USP14 inhibition has been considered a therapeutic strategy for accelerating degradation of aggregation-prone proteins in neurodegenerative diseases and for inhibiting proteasome function to induce apoptotic cell death in cancers. Here we studied the effects of USP14 inhibition in mammalian cells using small molecule inhibitors and an inactive USP14 mutant C114A. Neither the inhibitors nor USP14 C114A changed the level of TDP-43, tau or α-synuclein in HEK293T cells. However, USP14 C114A led to an accumulation of ubiquitinated proteins, which were isolated by ubiquitin immunoprecipitation and identified by mass spectrometry. Among these proteins we confirmed that ubiquitinated β-catenin was accumulated in the cells expressing USP14 C114A with biochemistry and molecular biology experiments. The proteasome binding of USP14 C114A is required for its effect on ubiquitinated proteins. UCHL5 is the other cysteine-protease deubiquitinase associated with the proteasome. Interestingly, the inactive mutant of UCHL5 C88A also caused an accumulation of ubiquitinated proteins in HEK293T cells but did not affect β-catenin. Using ubiquitin immunoprecipitation and mass spectrometry, we identified the accumulated ubiquitinated proteins in UCHL5 C88A expressing cells which are mostly distinct from those accumulated in USP14 C114A expressing cells. Among the identified proteins are well established proteasome substrates and proteasome subunits. Together our data suggest that USP14 and UCHL5 can deubiquitinate distinct substrates at the proteasome and regulate the ubiquitination of the proteasome itself which is tightly linked to its function.

## Introduction

The ubiquitin-proteasome system (UPS) is the main protein degradation pathway in eukaryotic cells (Collins and Goldberg, 2017). It determines the half-life of most cellular proteins, eliminates misfolded and damaged proteins, and is essential for protein homeostasis in cells. Proteins destined for degradation are tagged by the conjugation of a small 76-residue protein called ubiquitin, often in the form of polymeric chains, which enable the substrate to be recognized and degraded by the proteasome (Wilkinson et al., 1980; Komander and Rape, 2012; Saeki, 2017). The 26S proteasome is composed of a 20S core particle (CP) containing the proteolytic chamber, and one or two 19S regulatory particles (RP) critical for substrate recognition, deubiquitination, unfolding, and translocation (Kish-Trier and Hill, 2013; Tomko and Hochstrasser, 2013). Prior to and during substrate degradation, its ubiquitin tag must be removed, ensuring efficient substrate translocation into the proteolytic chamber and ubiquitin recycling in the cell (Finley et al., 2016; de Poot et al., 2017). The 19S RP harbors three deubiquitinating enzymes (DUBs) including two cysteine proteases, ubiquitin-specific protease 14 (USP14 in mammals or Ubp6 in yeast) and ubiquitin C-terminal hydrolase L5 (UCHL5, also known as UCH37), and a Zn^2+^-dependent metalloprotease RPN11 (Lam et al., 1997; Borodovsky et al., 2001; Leggett et al., 2002; Yao and Cohen, 2002).

RPN11 is essential to proteasomal degradation and loss of RPN11 activity causes accumulation of ubiquitinated proteins and cell death in budding yeast and mammalian cells (Verma et al., 2002; Yao and Cohen, 2002; Koulich et al., 2008; de Poot et al., 2017). No homolog of UCHL5 is present in the genome of budding yeast (Leggett et al., 2002). Loss of UCHL5 does not affect cell survival or change the level of free or conjugated ubiquitin (Koulich et al., 2008). USP14/Ubp6 is dynamically associated with the 19S RP through its ubiquitin-like (UBL) domain, not essential for cell survival, but important for ubiquitin recycling in budding yeast and a spontaneous ataxia mouse strain (Leggett et al., 2002; Chernova et al., 2003; Anderson et al., 2005). Loss of USP14 in mammalian cells does not alter free and conjugated ubiquitin levels but increases the degradation rate (Koulich et al., 2008; Kim and Goldberg, 2017).

It was reported that USP14-mediated deubiquitination may lead to release of the otherwise competent substrate from the proteasome and prevent its degradation, therefore inhibition of USP14 may accelerate degradation of such substrates (Hanna et al., 2006; Lee et al., 2010; Homma et al., 2015; Boselli et al., 2017). But this effect was not observed in other studies (Jin et al., 2012; Hyrskyluoto et al., 2014; Ortuno et al., 2016; Kiprowska et al., 2017). A small molecule USP14 inhibitor IU1 and an inactive USP14 mutant C114A were shown to reduce the level of the amyotrophic lateral sclerosis (ALS) and dementia disease proteins TDP-43 and tau in mammalian cells (Lee et al., 2010). Another small molecule inhibitor of USP14 and UCHL5, b- AP15, was shown to induce an accumulation of ubiquitinated proteins and apoptotic cell death in multiple myeloma cells (D’Arcy et al., 2011). Therefore, USP14 has been considered a drug target for neurodegenerative diseases and cancer although the former by enhancing and the latter by inhibiting proteasome function.

We were interested in USP14 as a potential drug target for neurodegenerative diseases. We started our study by repeating the experiments where USP14 WT or the inactive mutant C114A was co-expressed with TDP-43, tau or α-synuclein in HEK293T cells (Lee et al., 2010; Ortuno et al., 2016). Neither USP14 WT nor C114A changed the protein level of TDP-43, tau or α- synuclein which is consistent with the data reported by Miller and colleagues (Ortuno et al., 2016). Surprisingly, we observed a robust accumulation of high molecular weight K48-linked polyubiquitinated proteins with USP14 C114A expression. We reasoned these proteins are potential USP14 substrates, therefore combined immunoprecipitation and mass spectrometry analysis and identified a list of accumulated proteins including known proteasome substrates and multiple proteasome subunits. Among the identified proteins we focused on β-catenin and validated ubiquitinated β-catenin was indeed accumulated with USP14 C114A expression. These data prompted us to examine the effects of UCHL5 C88A in cells. Interestingly, UCHL5 C88A expression also caused an accumulation of high molecular weight K48-linked polyubiquitinated proteins but did not affect β-catenin. Our study supports that USP14 and UCHL5 can deubiquitinate specific substrates at the proteasome and are key factors of proteasome autoregulation by deubiquitination of proteasome subunits.

## Materials and Methods

### Plasmids and biochemical reagents

Plasmids containing myc-TDP-43 and myc-TDP-43 M337V were generated as described previously (Wobst et al., 2017). Plasmids for tau (4R/2N), α-synuclein, USP14 and UCHL5 were purchased from Origene. Mutant plasmids USP14 C114A, USP14 WT ΔUBL, USP14 C114A ΔUBL and UCHL5 C88A were generated from the wildtype plasmid by Genscript. Cells were treated with IU1 (25-100 µM; Selleckchem or Sigma), MG132 (10 µM; Selleckchem), PS341 (10 µM; Selleckchem) and b-AP15 (0.5-2 µM; Selleckchem). All compounds were dissolved in DMSO (Sigma), and control samples were treated with the same volume of DMSO.

### In vitro USP14 ubiquitin-rhodamine hydrolysis assay

Ubiquitin-Rhodamine 110 (Ub-Rho) and Vinyl sulphone-treated 26S-proteasomes (26S-VS) were purchased from Ubiquigent (60-0117-050 and 65-1020-BUL, respectively). Full length USP14 was purified as described previously (Lee et al., 2012). IU1 (Sigma) and b-AP15 (Millipore) were dissolved in DMSO and dispensed into black 384 well Greiner plates (Greiner Bio, 784076) over a gradient resulting in 100 µM to 3 nM final concentrations in the assay. 5 µL of 2X USP14/26S-VS solution was added to the assay plate in an assay buffer of 40 mM Tris-HCl pH 7.5, 0.01 % Brij-35, 1 mM TCEP. Reactions were started by adding 5 µL of 2X solution of Ub-Rho (2X concentration of 200 nM). Plates were covered and incubated at room temperature for 60 minutes, and the reactions were stopped by adding 5 µL of 100 mM citric acid. Plates were incubated for 15 minutes at room temperature and then read on a Tecan M1000 microplate reader (Fluorescence Intensity, Excitation wavelength: 485 nm, Excitation bandwidth: 10 nm, Emission wavelength: 520 nm, Emission bandwidth: 10 nm). IC_50_ values were determined by fitting curves to a log (compound concentration) vs. response - variable slope equation using GraphPad Prism 7.0.

### Cell culture and transfections

HEK293T cells (ATCC) were cultured in DMEM (Thermo Fisher) containing 10% FBS and 1% penicillin/streptomycin at 37°C with 5% CO2 in a humidified incubator. For transfections, cells were plated on 6-well plates or 10 cm plates 24 h prior to transfection. Cells were transfected with plasmids using Lipofectamine 2000 (Thermo Fisher) according to manufacturer’s instructions. Medium was changed 24 hours after transfection and cells were lysed 48 hours after transfection unless stated otherwise.

### Cell lysis and immunoblotting

Cells were scraped in an appropriate volume of RIPA buffer supplemented with a protease inhibitor cocktail (Complete Mini EDTA free tablets, Sigma-Aldrich). The cell lysate was centrifuged at 10,000 g for 10 minutes at 4 °C. The supernatant was collected, and protein concentrations were measured using the BCA assay (Pierce). 20-50 µg of total proteins were mixed with LDS buffer (Thermo Fisher) containing 20% β-mercaptoethanol and heated at 55°C for 10 minutes. The samples were centrifuged briefly and separated using 4-12% Bis-Tris gels or 4-12% Bolt gels (Thermo Fisher) and transferred onto polyvinylidene difluoride (PVDF) membranes (Millipore). The blots were blocked with 5% milk or 5% BSA in 1x TBST and probed with the following antibodies: α-synuclein (10841-1-AP, Proteintech, 1:1000), α-tubulin (mouse 7291, Abcam, 1:5000), α-tubulin (rabbit 52866, Abcam, 1:5000), β-catenin (610153, BD Transduction Laboratories, 1:2000), β-tubulin (T8328, Sigma, 1:5000), FK2 (PW8810, Enzo Life Sciences, 1:10000), GAPDH (sc-32233, Santa Cruz Biotechnology Inc., 1:10000), HA tag (3724, Cell Signaling Technology, 1:2000), tau (Monoclonal MN1000, Thermo Fisher, 1:1000), TDP-43 (10782-2-AP, Proteintech, 1:5000), TfR (Transferrin Receptor/CD71 Antibody #13-6800, Thermo Fisher, 1:1000), ubiquitin (3933, Cell Signaling Technology, 1:1000), K48-linked polyubiquitin (#05-1307, Millipore, 1:1000), K63-linked polyubiquitin (#05-1308, Millipore, 1:1000), UCHL5 (sc-271002, Santa Cruz Biotechnology Inc., 1:1000), USP14 (WH0009097M4, Sigma Aldrich, 1:1000). Blots were then incubated with mouse or rabbit secondary antibodies conjugated with HRP (Jackson Immunoresearch). Bands were detected using ECL substrate (Super Signal West Dura, Thermo Scientific) and imaged using the BioRad Universal Hood III Imager and BioRad Image Lab software. Bands were quantified using Quantity One (BioRad) and normalized to housekeeping genes in the same samples and then all samples were normalized to the average of control samples in the same experiments. We used GAPDH, α-tubulin, β-tubulin and TfR as housekeeping genes which did not change with experimental conditions.

### Immunoprecipitation

One hundred µg of proteins were mixed with RIPA buffer to a final volume of 500 µl. Protein A/G agarose beads (Pierce) were added to the above protein samples and rotated at 4 °C for 1 h for preclearing. The samples were then centrifuged at 8,000 g for 1 minute. 10% of the supernatant was used as input to confirm expression and loading. 2-10 µg of specific antibodies or control mouse or rabbit IgG antibodies (Jackson Immunoresearch) were added to the remaining supernatant and rotated at 4 °C for 1 h. Then fresh protein A/G agarose beads (Thermo Fisher) were added to the above samples and rotated at 4 °C overnight to pulldown antibody bound proteins. The samples were centrifuged at 8,000 g for 1 minute. The beads were washed with RIPA buffer four times and the immunoprecipitated complexes were detached from the beads by boiling the samples at 95 °C for 10 minutes in 30 µl of LDS loading buffer containing 5% β- mercaptoethanol. The samples were centrifuged at 10,000 g for 15 minutes and the supernatants were analyzed using immunoblotting.

### IP for mass spectrometry analysis

Scaled-up IP was performed as described above with 3700 µg of proteins. The K48 immunoprecipitates from USP14 WT, USP14 C114A, UCHL5 WT and UCHL5 C88A samples, and the control rabbit IgG immunoprecipitates from USP14 C114A and UCHL5 C88A samples were loaded on a 4-12% Bis-Tris gel (Thermo Fisher). The gel was stained using Coomassie blue (EZBlue Stain, Sigma) and destained using dH_2_O. The gel pieces above the heavy chain were cut and analyzed using mass spectrometry.

### Mass spectrometry

Trypsin digestion, liquid chromatography-tandem mass spectrometry (LC-MS/MS), and MS/MS spectral search in the human database (Uniprot) using the Sequest 28 analysis program was performed by Taplin Mass Spectrometry Facility (Harvard University).

To identify proteins specifically accumulated in USP14 C114A and UCHL5 C88A samples, proteins detected in the IgG control sample were removed without further consideration. The remaining proteins that were enriched using the Sum Intensity parameter in the C114A and C88A samples compared to the WT samples in all three replicates were included in Table 1, Table 2, Supplementary Table 1 and Supplementary Table 3. The lists of proteins in Table 1 and Table 2 were analyzed using Gene Ontology to identify pathways that were significantly enriched.

**Table 1.**
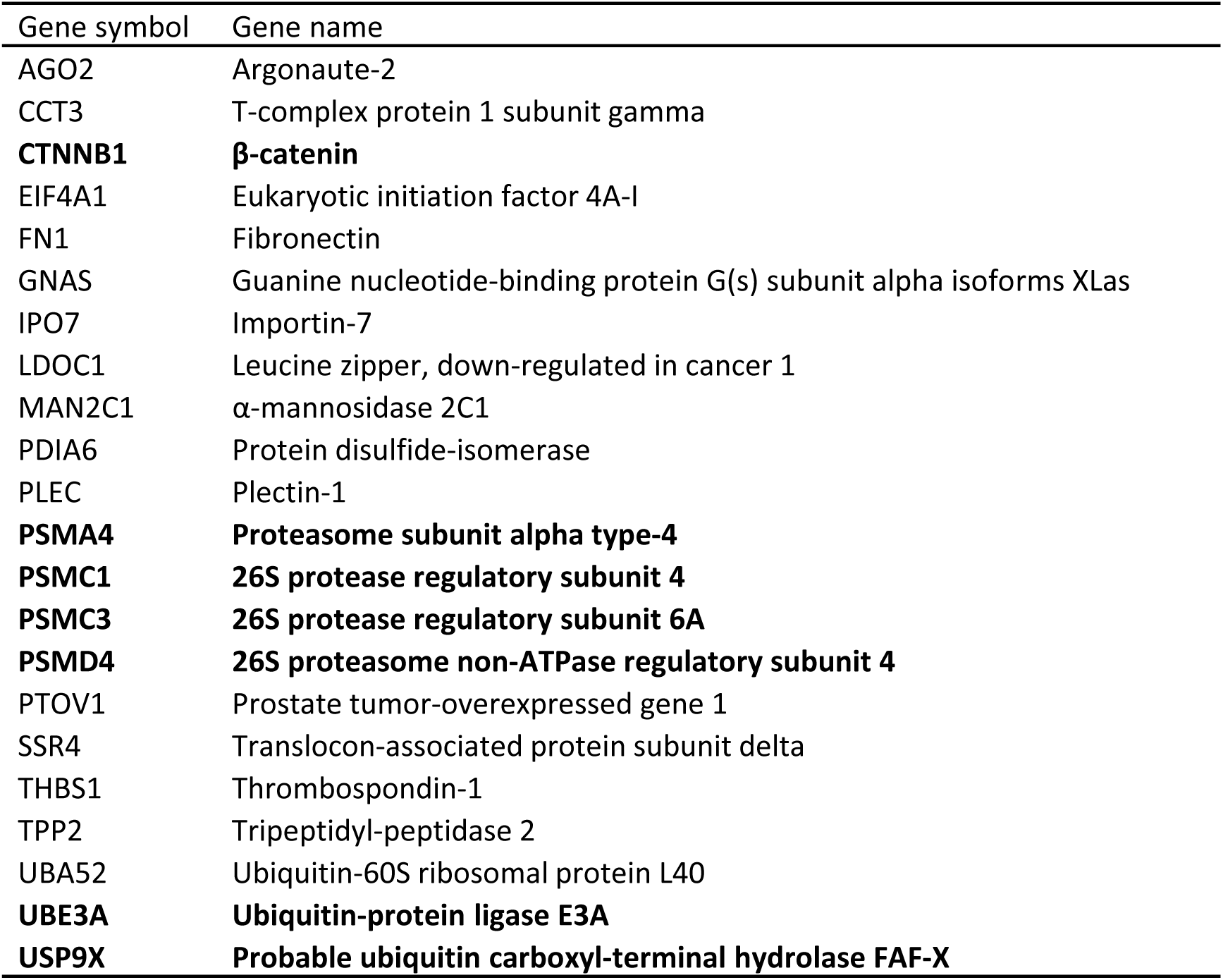
Ubiquitinated proteins accumulated in the cells expressing USP14 C114A.

**Table 2.** Ubiquitinated proteins accumulated in the cells expressing UCHL5 C88A.

| Gene symbol | Gene name |
| --- | --- |
| AEBP1 | Adipocyte enhancer-binding protein 1 |
| ASNS | Asparagine synthetase |
| ATXN10 | Ataxin-10 |
| CALR | Calreticulin |
| CCT3 | T-complex protein 1 subunit gamma |
| CEP135 | Centrosomal protein of 135 kDa |
| COPB1 | Coatamer subunit beta |
| CUL3 | Cullin 3 |
| DNAJC10 | DnaJ homolog subfamily C member 10 |
| FN1 | Fibronectin |
| HYOU1 | Hypoxia up-regulated protein 1 |
| IPO5 | Importin-5 |
| IPO9 | Importin-9 |
| MAMDC2 | MAM domain-containing protein 2 |
| NACA | Nascent polypeptide-associated complex subunit alpha |
| NCKAP1 | Nck-associated protein 1 |
| NID1 | Nidogen-1 |
| PDIA6 | Protein disulfide-isomerase A6 |
| PLOD1 | Procollagen-lysine,2-oxoglutarate 5-dioxygenase 1 |
| **PSMA4** | **Proteasome subunit alpha type-4** |
| **PSMC3** | **26S protease regulatory subunit 6A** |
| **PSMD1** | **26S proteasome non-ATPase regulatory subunit 1** |
| RPS3 | 40S ribosomal protein S3 |
| THBS1 | Thrombospondin-1 |
| THOP1 | Thimet oligopeptidase |
| **UCHL5** | **Ubiquitin carboxyl-terminal hydrolase isozyme L5** |
| UGGT1 | UDP-glucose:glycoprotein glucosyltransferase 1 |
| VARS | Valine-tRNA ligase |

### Statistics

GraphPad Prism 7.0 was used for statistical analysis. One-way analysis of variance (ANOVA) was used to test for statistical significance in all figures except figure 7 where t-test was used to determine statistical significance. Values are mean ± standard deviation (SD). Statistical significance was set at *P* < 0.05.

## Results

### Inactive USP14 C114A does not change the level of TDP-43, tau or α-synuclein in HEK293T cells

Previous studies (Lee et al., 2010; Ortuno et al., 2016) have reported findings regarding effects of USP14 overexpression or inhibition on degradation of aggregation-prone proteins in neurodegenerative diseases. We repeated the experiments by co-expressing the empty vector control, USP14 WT or C114A with myc-TDP-43, tau or α-synuclein in HEK293T cells (Fig. 1A and B). Neither USP14 WT nor C114A changed the protein level of endogenous TDP-43, myc-TDP-43 WT, myc-TDP-43 M337V (an ALS causal mutation, (Sreedharan et al., 2008)), tau or α-synuclein.

**Figure 1.**
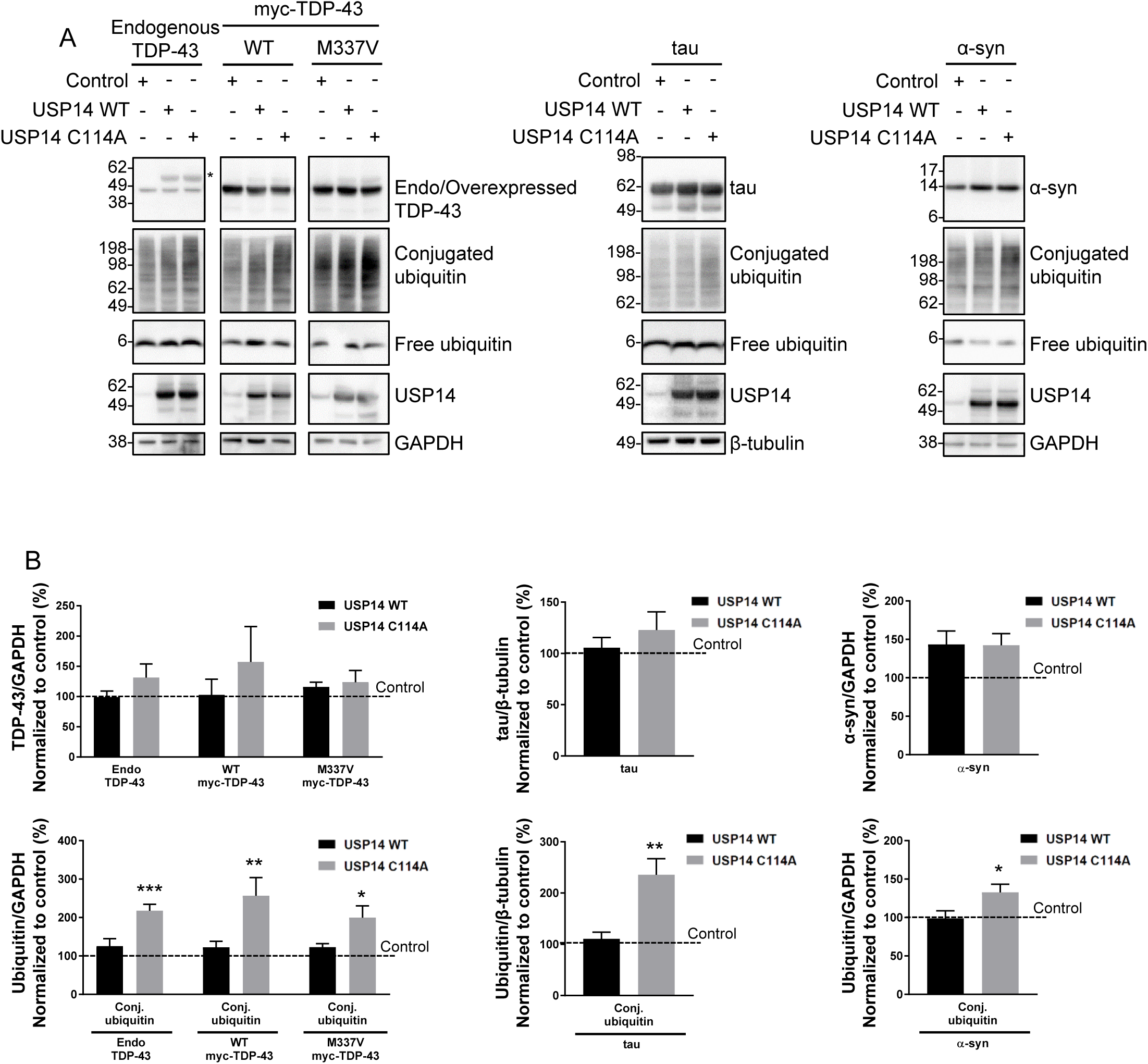
Expression of USP14 C114A has no effect on TDP-43, tau or α-synuclein but leads to an accumulation of ubiquitinated proteins. (A) Immunoblot showing the effects of a control plasmid, USP14 WT or C114A co-expression with myc-TDP-43 WT, myc-TDP-43 M337V, tau or α-synuclein in HEK293T cells. * Non-specific band. (B) USP14 WT or C114A has no effect on the protein level of TDP-43, tau and α-synuclein. Quantification of protein levels from A. n = 3-6, error bars represent SEM. (C) USP14 C114A causes an accumulation of polyubiquitinated proteins. Quantification of conjugated ubiquitin (conj. ubiquitin) from A. n = 3-6, error bars represent SEM, * *P* < 0.05, ** *P* < 0.01, *** *P* < 0.001.

We also tested two published USP14 inhibitors, IU1 (Lee et al., 2010) and b-AP15 (D’Arcy et al., 2011) in the ubiquitin-rhodamine hydrolysis assay with purified USP14 in the presence of vinyl sulphone-treated 26S proteasome (Hassiepen et al., 2007) (Supplementary Fig. 1). IU1 inhibited USP14 with an IC_50_ of 5.51 μM as previously published (Lee et al., 2010) whereas b-AP15 did not show significant inhibition up to 100 μM, likely due to that the proteasome was treated with vinyl sulphone (Supplementary Fig. 1). IU1 treatment (75 μM, 6 hours) did not affect the level of TDP-43, tau or α-synuclein expressed in HEK293T cells (Supplementary Fig. 2A and B). Proteasome inhibitors (MG132 10 µM + PS341 10 µM, 6 hours) as expected caused a robust accumulation of ubiquitinated proteins but did not affect the level of TDP-43, tau or α-synuclein (Supplementary Fig. 2A and B). Dose responses of 1U1 up to 100 μM (6 hours) also did not affect the level of TDP-43, tau or α-synuclein (Supplementary Fig. 2C and D). b-AP15 treatment caused an accumulation of ubiquitinated proteins as previously published (D’Arcy et al., 2011) but did not affect the level of TDP-43, tau or α-synuclein (Supplementary Fig. 2E and F).

**Fig. 2.**
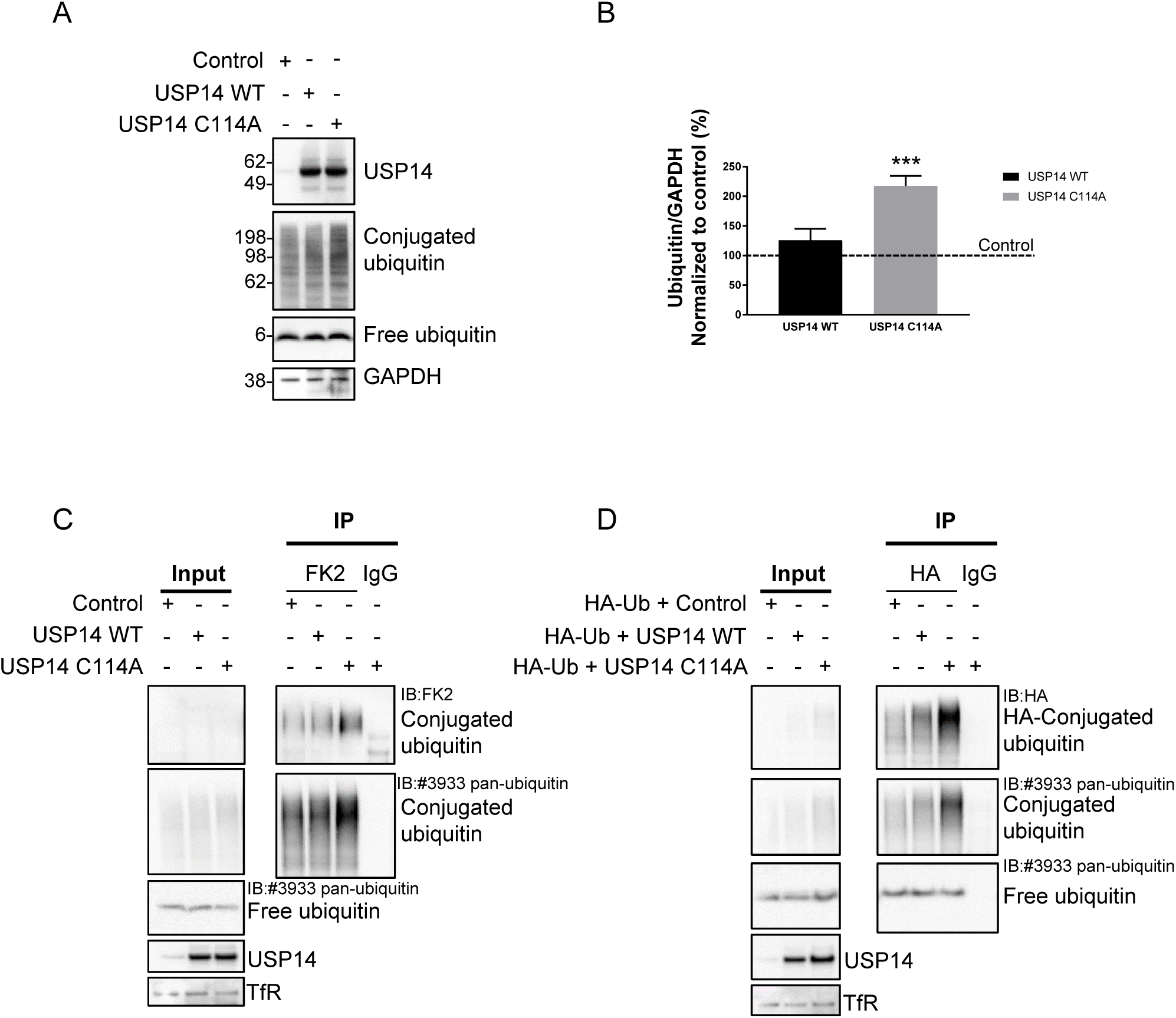
Inactive USP14 C114A causes accumulation of ubiquitinated proteins in HEK293T cells. (A) Immunoblot showing expression of USP14 C114A in HEK293T cells causes an accumulation of ubiquitinated proteins. (B) Quantification of conjugated ubiquitin from (A). *** *P* < 0.001, n = 4, error bars represent SEM. (C) Ubiquitin IP with the FK2 antibody from the cells expressing the indicated proteins, followed by IB with FK2 and the pan-ubiquitin antibody #3933. (D) IP using the HA antibody from the cells co-expressing HA-ubiquitin with the indicated proteins, followed by IB with the HA and pan-ubiquitin antibodies.

Although neither USP14 C114A nor USP14 inhibitors affected the levels of TDP-43, tau or α-synuclein, USP14 C114A expression consistently caused accumulation of ubiquitinated proteins in HEK293T cells in all the experiments (Fig. 1).

### Inactive USP14 C114A causes accumulation of ubiquitinated proteins in HEK293T cells

We next expressed the empty vector, USP14 WT or C114A in HEK293T cells to understand whether USP14 C114A has the same effect on ubiquitinated proteins in the absence of co-expressed TDP-43, tau or α-synuclein. USP14 C114A alone caused a similar accumulation of ubiquitinated proteins (Fig. 2A and B). Using an immunoprecipitation compatible ubiquitin antibody FK2, we were able to pull down more ubiquitin conjugates from the cells expressing USP14 C114A compared to the empty vector control or WT (Fig. 2C). Furthermore, using an HA antibody, we were able to pull down more ubiquitin conjugates from the cells co-transfected with HA-ubiquitin and USP14 C114A compared to the cells co-transfected with the empty vector control or WT (Fig. 2D). These data confirm inactive USP14 causes ubiquitinated protein accumulation in cells.

### Identification of the accumulated ubiquitinated proteins induced by USP14 C114A

We next assessed the chain linkage of the ubiquitin conjugates accumulated with USP14 C114A expression. We focused on lysine 48 (K48)- and lysine 63 (K63)-linked chains as these are the main ubiquitin tags known to target substrates for proteasomal degradation (Glickman and Ciechanover, 2002; Jacobson et al., 2009). K48-linked but not K63-linked ubiquitin conjugates were accumulated in the cells expressing USP14 C114A (Fig. 3A and B). To identify these proteins, we combined immunoprecipitation with the antibody against K48-linked ubiquitin chains (simplified as K48 antibody hereafter) and mass spectrometry (IP-MS). K48-linked ubiquitinated proteins were immunoprecipitated from the cells expressing USP14 WT or C114A and separated by SDS-PAGE, followed by immunoblotting (IB) (Fig. 3C) or staining with Coomassie blue (Fig. 3D), both of which showed more ubiquitinated proteins were pulled down from the USP14 C114A than WT samples. We also included a control IgG IP from USP14 C114A expressing cells to be able to filter out proteins that were pulled down non-specifically (Fig. 3C and D). The lanes were cut above the antibody heavy chain and subjected to MS analysis (Fig. 3D). The IP-MS was repeated in triplicate and all the proteins identified in the IgG IP were removed without further consideration. The remaining proteins more abundant in the USP14 C114A than WT samples in all three replicates are listed in Table 1 (additional details in Supplementary Table 1 and 2).

**Fig. 3.**
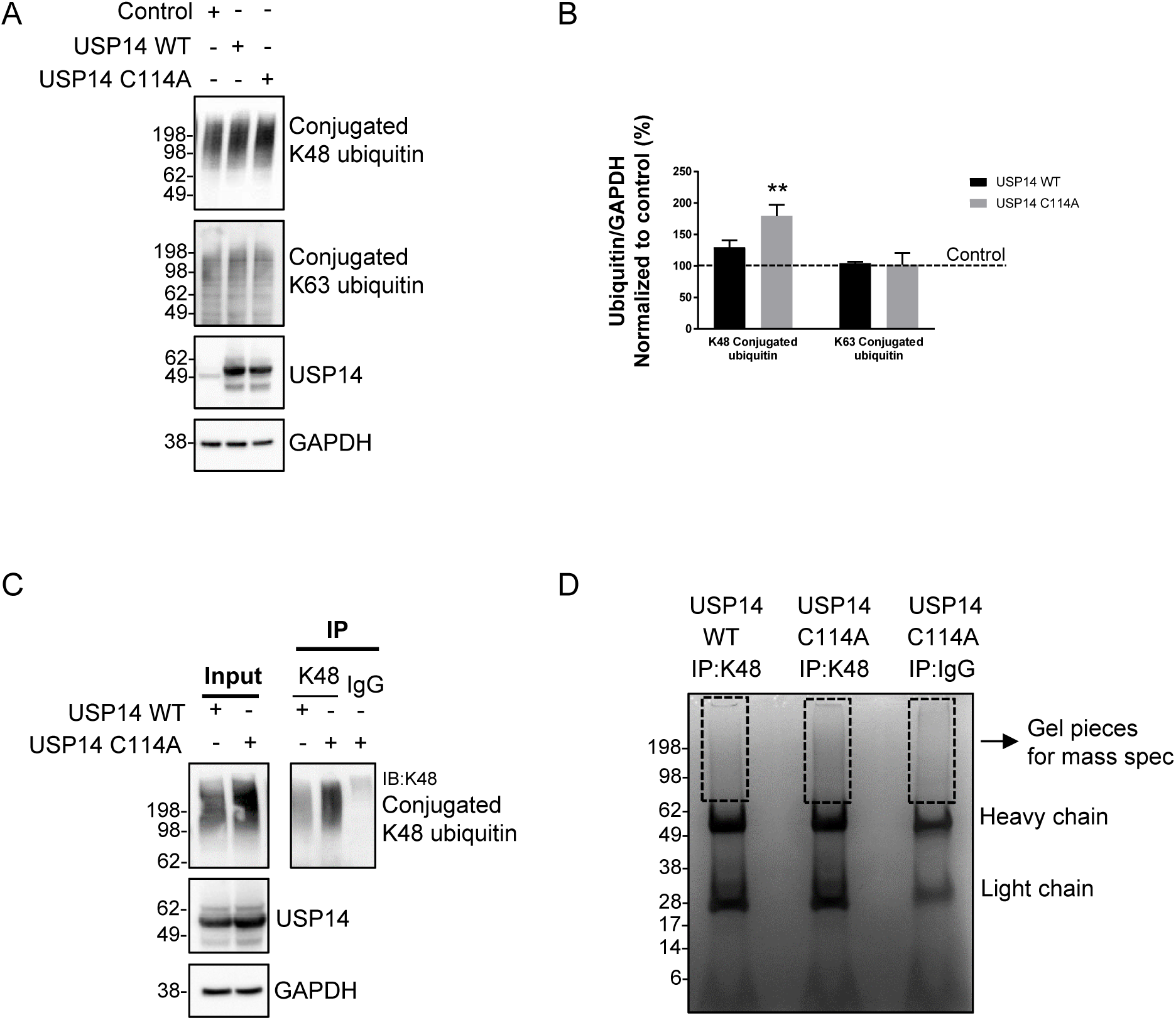
Isolation of the ubiquitinated proteins accumulated in the cells expressing USP14 C114A. (A) Immunoblot showing K48-linked but not K63-linked ubiquitinated proteins are accumulated in the cells expressing USP14 C114A. (B) Quantification of protein levels from (A). n = 3, ** *P* < 0.01, error bars represent SEM. (C) and (D) K48-linked ubiquitinated proteins were immunoprecipitated from the cells expressing the indicated proteins. A control IP was done with IgG from the cells expressing USP14 C114A. The IP samples were subjected to (C) IB with the K48 antibody or (D) Coomassie blue staining. The gel pieces above the antibody heavy chain in (D) were cut for mass spectrometry analysis.

Among the list are well-established proteasomal substrates and important signaling molecules such as β-catenin (Stamos and Weis, 2013) and Argonaute-2 (Smibert et al., 2013) (Table 1). Multiple proteasome subunits were identified including PSMA4 (20S subunit α3), PSMC1 (19S AAA-ATPase subunit Rpt2), PSMC3 (19S AAA-ATPase subunit Rpt5), and PSMD4 (19S non-ATPase subunit Rpn10/S5A). The Angelman syndrome disease gene ubiquitin-protein ligase E3A (UBE3A) (Sell and Margolis, 2015) and mental retardation disease gene ubiquitin specific peptidase, X-linked (USP9X) (Homan et al., 2014) were also identified. The IP-MS data suggest USP14 regulates not only substrate deubiquitination but also the ubiquitination status of the proteasome.

### Confirmation that ubiquitinated β-catenin is accumulated in USP14 C114A expressing cells

We focused on β-catenin to validate the IP-MS data. A higher molecular weight smear of the β- catenin band was observed in the USP14 C114A samples, suggesting an accumulation of ubiquitinated β-catenin (Fig. 4A). The total protein level of β-catenin was increased in the cells expressing USP14 C114A compared to the empty vector control or WT (Fig. 4A and B). The level of ubiquitin conjugates was also significantly increased as seen previously (Fig. 2, 4A and 4B).

**Fig. 4.**
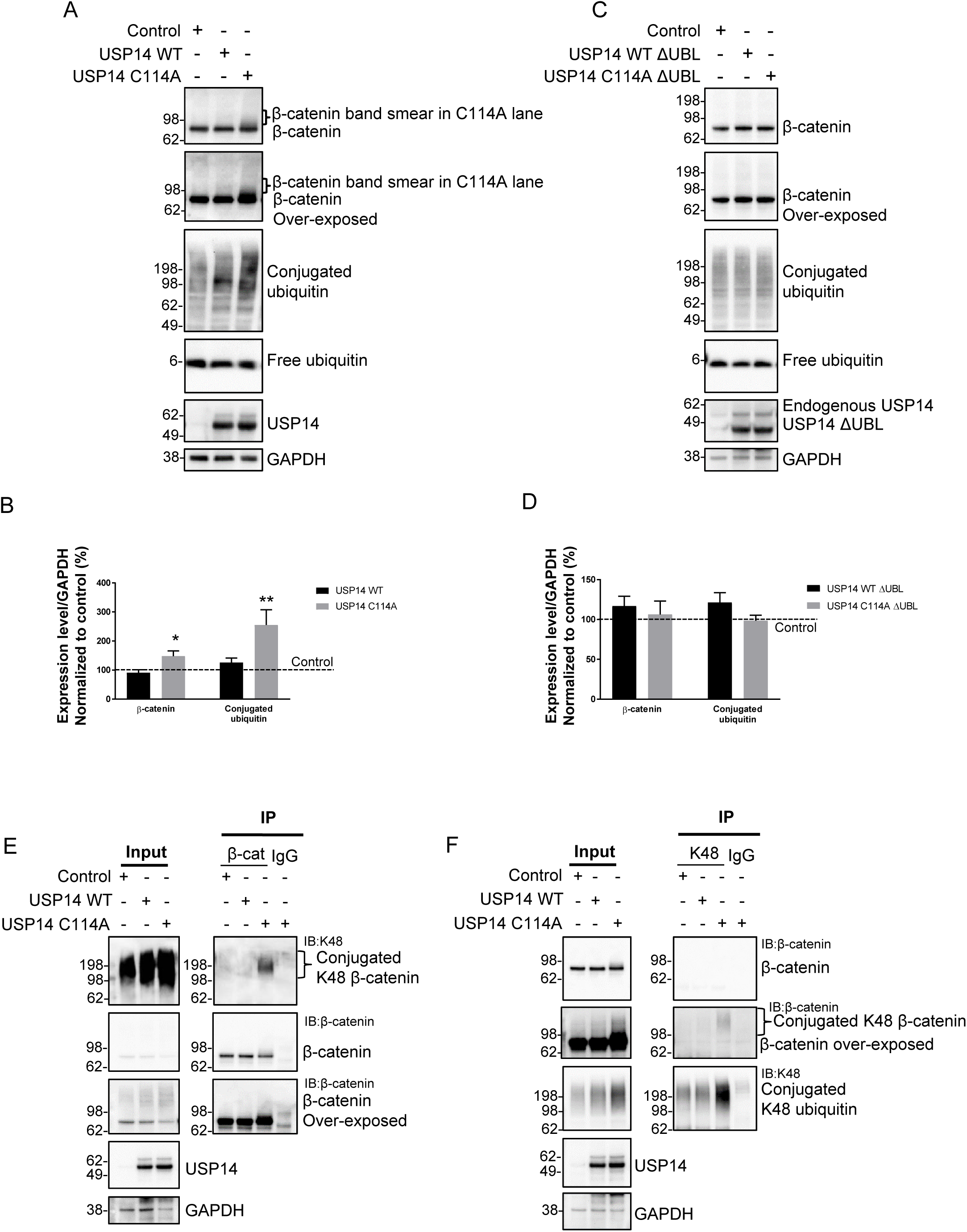
Confirmation that ubiquitinated β-catenin is accumulated in USP14 C114A expressing cells. (A) Immunoblot showing USP14 C114A increases the total protein level of β-catenin in cells. Note the smearing of the β-catenin band in the C114A lane. (B) Quantification from (A). n = 6, * *P* < 0.05, ** *P* < 0.01, error bars represent SEM. (C) Immunoblot showing proteasome binding is required for the effects of USP14 C114A on β-catenin and conjugated ubiquitin. Deletion of the UBL proteasome binding domain (ΔUBL) abolishes the effects of USP14 C114A on β-catenin and conjugated ubiquitin. (D) Quantification from (C). n = 6, error bars represent SEM. (E) β-catenin IP followed by K48 IB showing ubiquitinated β-catenin is accumulated in the cells expressing USP14 C114A. (F) K48 IP followed by β-catenin IB showing ubiquitinated β-catenin is accumulated in the cells expressing USP14 C114A.

To confirm the effects of USP14 C114A are dependent on proteasome binding, we generated USP14 WT and C114A constructs lacking the UBL domain, USP14 WT ΔUBL and C114A ΔUBL. The ΔUBL constructs were predicted to be ~10 kDa smaller than the full-length constructs and were expressed as expected (Fig. 4C). USP14 C114A ΔUBL had no effect on the levels of β- catenin or ubiquitin conjugates, demonstrating that proteasome binding is required for USP14 C114A’s effect on ubiquitinated proteins (Fig. 4C and D).

To further confirm the accumulation of ubiquitinated β-catenin in the cells expressing USP14 C114A, we conducted mutual IP-IB experiments using β-catenin or K48 antibodies (Fig. 4E and F). β-catenin IP showed a smear of K48-linked ubiquitinated β-catenin from USP14 C114A expressing cells (Fig. 4E). The reverse approach of K48 IP and β-catenin IB also showed a high molecular weight smear of ubiquitinated β-catenin from USP14 C114A expressing cells (Fig. 4F). Taken together, these data confirm that ubiquitinated β-catenin is accumulated in USP14 expressing cells.

### Inactive UCHL5 C88A causes accumulation of ubiquitinated proteins but does not affect β- catenin

To understand whether USP14 has a specific effect on β-catenin or the other proteasomal cysteine-protease deubiquitinase UCHL5 also shares the same effect, we generated a catalytically inactive mutant of UCHL5 by mutating the catalytic cysteine at position 88 to alanine (UCHL5 C88A). This mutant was shown to lack deubiquitinase activity but bind to the proteasome similarly to WT (Yao et al., 2006). Interestingly, expression of the inactive UCHL5 C88A also led to an accumulation of K48-linked ubiquitin conjugates (Fig. 5A and B). Furthermore UCHL5 C88A expression led to a high molecular weight laddering of UCHL5 itself as detected by IB with a UCHL5 antibody, presumably representing ubiquitinated UCHL5 (Fig. 5A) (Besche et al., 2014; Jacobson et al., 2014; Randles et al., 2016). While the IP-MS data suggest USP14 is required for deubiquitination of multiple proteasomal subunits (Table 1), USP14 C114A did not affect UCHL5 ubiquitination as shown by the lack of the UCHL5 laddering (Fig. 5C).

**Fig. 5.**
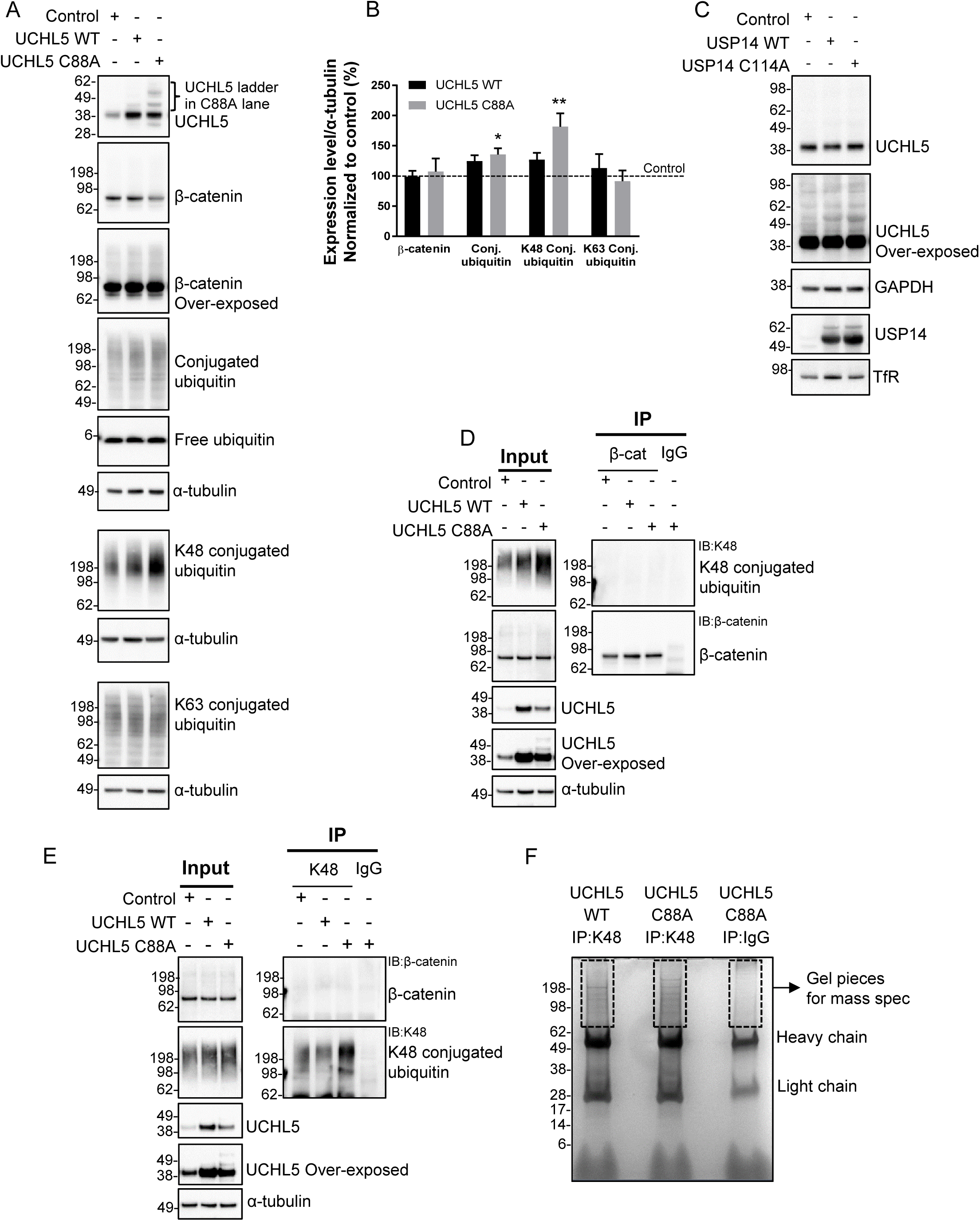
UCHL5 C88A causes an accumulation of ubiquitinated proteins but does not affect β- catenin degradation. (A) Immunoblot showing the effects UCHL5 WT and C88A in HEK293T cells. Note the lack of band smearing of β-catenin but the presence of band laddering of UCHL5 and the increase of conjugated ubiquitin in the C88A lane. (B) Quantification from (A). n = 3-6, * *P* < 0.05, ** *P* < 0.01, error bars represent SEM. (C) Immunoblot showing USP14 C114A does not induce UCHL5 band laddering in cells. (D) β-catenin IP followed by K48 IB showing the lack of ubiquitinated β-catenin accumulation in the cells expressing UCHL5 C88A. (E) K48 IP followed by K48 and β-catenin IB showing although the accumulated ubiquitinated proteins were immunoprecipitated but ubiquitinated β-catenin was not accumulated in the cells expressing UCHL5 C88A. (F) K48-linked ubiquitinated proteins were immunoprecipitated from the cells expressing the indicated proteins. A control IP was done with IgG from the cells expressing UCHL5 C88A. The IP samples were subjected to Coomassie blue staining. The gel pieces above the antibody heavy chain were cut for mass spectrometry analysis.

While UCHL5 C88A showed clear effects on ubiquitin conjugates and its own ubiquitination, it had no effect on β-catenin accumulation or band smearing (Fig. 5A and B). To rule out the possibility that the accumulation of ubiquitinated β-catenin was below the detection limit of straight IB, we conducted mutual IP-IB experiments using β-catenin and K48 antibodies, similar to those done with USP14 (Fig. 4E, 4F, 5D, and 5E). β-catenin IP failed to pull down any K48-linked ubiquitinated β-catenin from UCHL5 C88A expressing cells although the input samples confirmed the accumulation of K48-linked ubiquitin conjugates and UCHL5 laddering in those cells (Fig. 5D). K48 IP pulled down the accumulated ubiquitinated proteins from UCHL5 C88A expressing cells but β-catenin was not detected (Fig. 5E). These data indicate inactive USP14 but not inactive UCHL5 specifically affects β-catenin ubiquitination, suggesting β-catenin could be a specific substrate of USP14.

### Identification of the accumulated ubiquitinated proteins induced by UCHL5 C88A

To identify the accumulated ubiquitinated proteins induced by UCHL5 C88A, K48-linked ubiquitinated proteins were immunoprecipitated from UCHL5 WT or C88A expressing cells and subjected to SDS-PAGE followed by Coomassie blue staining for MS (Fig. 5F). Similarly to USP14, we included a control IgG IP from UCHL5 C88A expressing cells to filter out non-specific proteins. The IP-MS was repeated in triplicate and all the proteins identified in the IgG IP were removed without further consideration. The remaining proteins more abundant in UCHL5 C88A than WT samples in all three replicates are listed in Table 2 (additional details in Supplementary Table 3 and 4).

16 unique proteins were identified for USP14 and 22 for UCHL5, and 6 common proteins for both (Table 1 and 2, Fig. 6A). Importantly, ubiquitinated β-catenin was not enriched in UCHL5 C88A samples, which is consistent with the biochemical characterization (Fig. 4 and 5). Moreover, ubiquitinated UCHL5 was indeed enriched in UCHL5 C88A samples, which confirmed that the high molecular weight laddering of UCHL5 represented ubiquitinated species (Fig. 5A). Previously known proteasome substrates which are also disease genes were identified for UCHL5 such as Ataxin-10 (Li et al., 2011) and calreticulin (Goitea and Hallak, 2015). Ubiquitinated proteasome subunits PSMA4 and PSMC3 (Rpt5) were accumulated in both USP14 C114A and UCHL5 C88A samples, while ubiquitinated PSMC1 (Rpt2) and PSMD4 (Rpn10/S5A) were specific for USP14 and PSMD1 (Rpn2) was specific for UCHL5 (Fig. 6A). Taken together our biochemistry and IP-MS data suggest that USP14 and UCHL5 can deubiquitinate specific substrates at the proteasome.

**Fig. 6.**
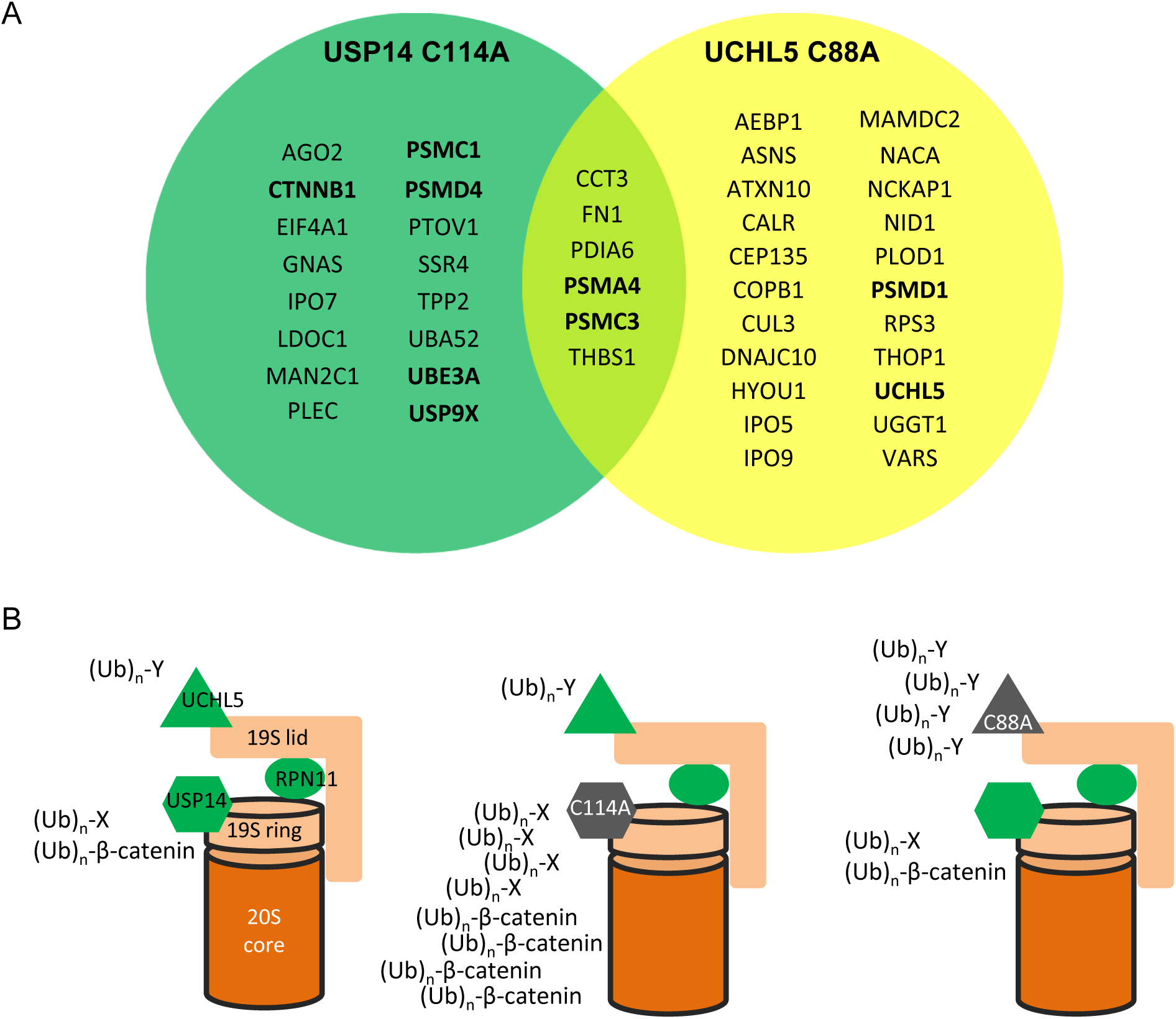
A model showing the effects of inactive USP14 and inactive UCHL5 at the proteasome. (A) Venn diagram of the ubiquitinated proteins accumulated in USP14 C114A and UCHL5 C88A expressing cells. (B) Inactive USP14 C114A causes accumulation of specific substrates (Ub)_n_-X including ubiquitinated β-catenin. Inactive UCHL5 C88A causes accumulation of specific substrates (Ub)_n_-Y.

## Discussion

In the absence of a ubiquitinated substrate, USP14/Ubp6 reduces basal peptidase and ATPase activity of the proteasome to degrade unstructured proteins lacking ubiquitination (Kim and Goldberg, 2017), supporting its function as a gate to prevent non-specific degradation and to allow entry of proper substrates. Ubiquitin loaded USP14 activates proteasomal ATPase, causes 20S gate opening, but inhibits RPN11, and delayed degradation of ubiquitinated substrates (Hanna et al., 2006; Peth et al., 2009; Peth et al., 2013; Bashore et al., 2015; Kim and Goldberg, 2017), supporting that it ensures sequential deubiquitination by itself first and then RPN11, and coordinates substrate unfolding and entry into the 20S CP. When loaded with a ubiquitinated substrate, UCHL5 also activates ATPase activity and 20S gate opening of the proteasome purified from USP14 knockout mammalian cells (Peth et al., 2013), suggesting UCHL5 may function as another gate to coordinate substrate entry and processing. USP14 and UCHL5 associate with the proteasome through different ubiquitin receptors, USP14 to Rpn1 at the base of the 19S RP and UCHL5 to Rpn13 at the top of the 19S RP (Collins and Goldberg, 2017) (Fig. 6B). Both USP14 and UCHL5 play complex catalytic and allosteric roles during substrate degradation by the proteasome.

In this study we identified that the inactive mutants USP14 C114A and UCHL5 C88A caused accumulations of distinct sets of ubiquitinated proteins in HEK293T cells (Fig. 1, 2, 3, 5, and 6). Both mutants retain the activity to bind to the proteasome (Yao et al., 2006; Lee et al., 2011). The UBL proteasome-binding domain of USP14 is required for C114A’s effect on ubiquitinated proteins (Fig. 4C). Among the accumulated proteins identified for USP14 C114A, we focused on β-catenin and confirmed its accumulation in the cells expressing USP14 C114A but not in the cells expressing UCHL5 C88A (Fig. 4 and 5). Besides β-catenin, Argonaute-2 (Fig. 6A, Table 1 and 2) was identified specifically for USP14 and is also a known proteasome substrate with important regulatory functions in small RNA-mediated silencing (Smibert et al., 2013). Previously known proteasomal substrates have also been identified specifically for UCHL5 such as Ataxin-10 (Li et al., 2011) and calreticulin (Goitea and Hallak, 2015) (Fig. 6A, Table 1 and 2). Interestingly, our IP-MS analysis show USP14 C114A causes the accumulation of ubiquitinated UBE3A and proteasome subunits PSMA4 (20S subunit α3), PSMC1 (19S AAA-ATPase subunit Rpt2), PSMC3 (19S AAA-ATPase subunit Rpt5), and PSMD4 (19S non-ATPase subunit Rpn10/S5A) (Fig. 6A). UCHL5 C88A causes the accumulation of ubiquitinated PSMA4, PSMC3, and PSMD1 (Rpn2) and is required for its own deubiquitination (Fig. 5 and 6A). Ubiquitination of Rpt5, Rpn10/S5A, Rpn13/Adrm1, and UCHL5 is induced by proteasome-associated UBE3A and UBE3C at the purified proteasome (Jacobson et al., 2014). Ubiquitin aldehyde further increases ubiquitination of these proteins, demonstrating USP14 and UCHL5 antagonize UBE3A and UBE3C to deubiquitinate these proteins (Jacobson et al., 2014). Ubiquitination of proteasome subunits impairs substrate recognition and processing by the proteasome and cellular stresses can modulate the ubiquitination status of the proteasome (Besche et al., 2014; Jacobson et al., 2014). Together our data suggest that USP14 and UCHL5 can deubiquitinate specific substrates at the proteasome and regulate the proteasome activity by modulating the ubiquitination status of the proteasome itself.

A recent study, combining USP14 interactome and quantitative proteomics in response to USP14 knockdown, identified fatty acid synthase (FASN) as a specific USP14 substrate, and suggested a list of proteins as potential USP14 substrates (Liu et al., 2018). All these studies including ours start to reveal a picture of USP14 substrates. In future studies, it is important to understand whether specific proteasomal substrates require USP14 and/or UCHL5 for degradation. It is also important to distinguish the roles of USP14 in degradation inhibition by removing the ubiquitin chains of substrates and in degradation facilitation by deubiquitinating substrates to allow unfolding.

We were interested in the effect of USP14 inhibition on reducing neurodegenerative disease proteins such as α-synuclein, TDP-43, and tau, but neither USP14 inhibitors nor USP14 C114A reduced the level of these proteins in HEK293T cells (Fig. 1 and Supplementary Fig. 2). Similar findings were also reported by Miller and colleagues (Ortuno et al., 2016). TDP-43 and α- synuclein have long half-lives and it is demonstrated that autophagy plays an important role in their degradation, and both proteasome and autophagy appear to be important for tau degradation (Cuervo et al., 2004; Chesser et al., 2013; Barmada et al., 2014). Treating cells with proteasome inhibitors did not modulate the levels of TDP-43, tau or α-synuclein (Fig. 1A and Supplementary Fig. 1A), suggesting these proteins are poor substrates of the proteasome in HEK293T cells, which could explain the lack of effect of USP14 inhibition on degradation of these proteins. Further studies are required to demonstrate whether USP14 inhibition indeed reduces the level of these proteins in neurons.

## Supporting information

## Acknowledgements

We thank Drs. Lin Song and Celina D’Cruz for their helpful advice during this study.

## Author Contributions

Participated in research design: Chadchankar, Doig, Jacobsen, Brandon, Moss, Wang

Conducted experiments: Chadchankar, Korboukh

Contributed new reagents or analytic tools: Chadchankar

Performed data analysis: Chadchankar, Korboukh, Wang

Wrote or contributed to the writing of the manuscript: Chadchankar, Wang

## Footnotes

This work was supported by funding from AstraZeneca. SJM is supported by National Institutes of Health (NIH)–National Institute of Neurological Disorders and Stroke Grants NS051195, NS056359, NS081735, R21NS080064 and NS087662; NIH–National Institute of Mental Health Grant MH097446, and DOD, AR140209.

---

**Supplementary Fig. 1.**
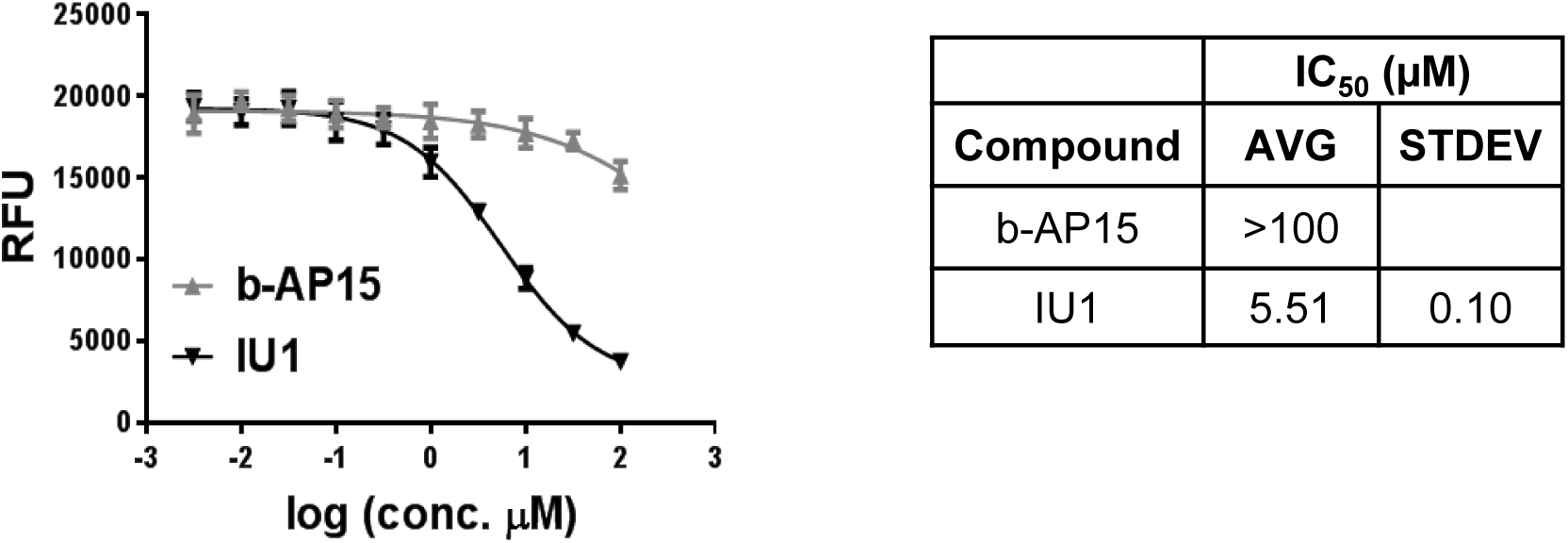
The activity of 1U1 and b-AP15 on USP14 in the ubiquitin-rhodamine hydrolysis assay.

**Supplementary Fig. 2.**
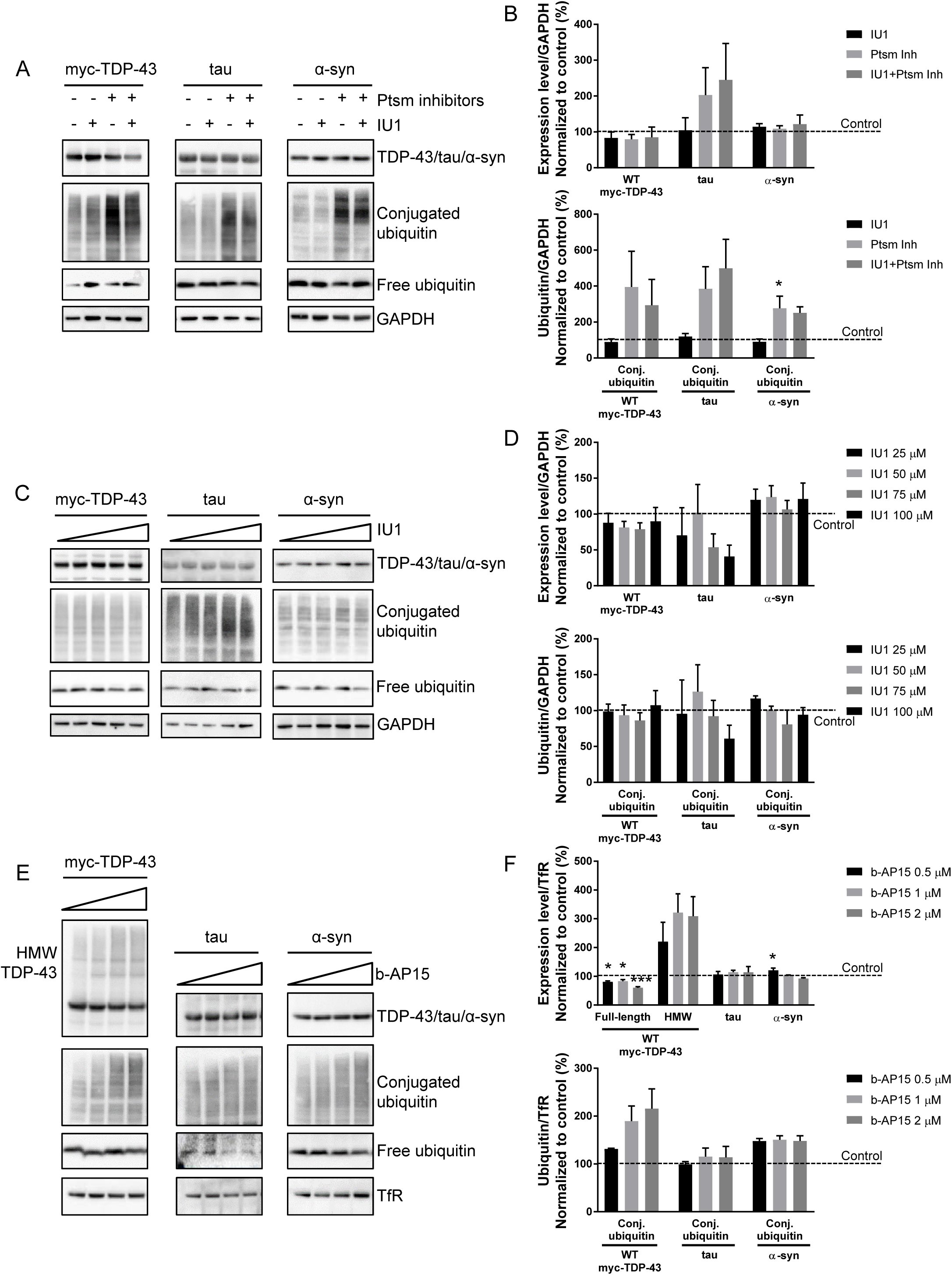
USP14 inhibition using the small molecule inhibitors IU1 or b-AP15 does not change the levels of TDP-43, tau or α-synuclein in HEK293T cells.

**Supplementary Table 1.** Quantification of polyubiquitinated proteins more abundant in the USP14 C114A than WT samples in all three replicates.

**Supplementary Table 2.** GO analysis of polyubiquitinated proteins more abundant in the USP14 C114A than WT samples in all three replicates.

**Supplementary Table 3.** Quantification of polyubiquitinated proteins more abundant in the UCHL5 C88A than WT samples in all three replicates.

**Supplementary Table 4.** GO analysis of polyubiquitinated proteins more abundant in the UCHL5 C88A than WT samples in all three replicates.

## References

Anderson C, Crimmins S, Wilson JA, Korbel GA, Ploegh HL and Wilson SM (2005) Loss of Usp14 results in reduced levels of ubiquitin in ataxia mice. J Neurochem 95:724–731.

Barmada SJ, Serio A, Arjun A, Bilican B, Daub A, Ando DM, Tsvetkov A, Pleiss M, Li X, Peisach D, Shaw C, Chandran S and Finkbeiner S (2014) Autophagy induction enhances TDP43 turnover and survival in neuronal ALS models. Nat Chem Biol 10:677–685.

Bashore C, Dambacher CM, Goodall EA, Matyskiela ME, Lander GC and Martin A (2015) Ubp6 deubiquitinase controls conformational dynamics and substrate degradation of the 26S proteasome. Nat Struct Mol Biol 22:712–719.

Besche HC, Sha Z, Kukushkin NV, Peth A, Hock EM, Kim W, Gygi S, Gutierrez JA, Liao H, Dick L and Goldberg AL (2014) Autoubiquitination of the 26S proteasome on Rpn13 regulates breakdown of ubiquitin conjugates. EMBO J 33:1159–1176.

Borodovsky A, Kessler BM, Casagrande R, Overkleeft HS, Wilkinson KD and Ploegh HL (2001) A novel active site-directed probe specific for deubiquitylating enzymes reveals proteasome association of USP14. EMBO J 20:5187–5196.

Boselli M, Lee BH, Robert J, Prado MA, Min SW, Cheng C, Silva MC, Seong C, Elsasser S, Hatle KM, Gahman TC, Gygi SP, Haggarty SJ, Gan L, King RW and Finley D (2017) An inhibitor of the proteasomal deubiquitinating enzyme USP14 induces tau elimination in cultured neurons. J Biol Chem.

Chernova TA, Allen KD, Wesoloski LM, Shanks JR, Chernoff YO and Wilkinson KD (2003) Pleiotropic effects of Ubp6 loss on drug sensitivities and yeast prion are due to depletion of the free ubiquitin pool. J Biol Chem 278:52102–52115.

Chesser AS, Pritchard SM and Johnson GV (2013) Tau clearance mechanisms and their possible role in the pathogenesis of Alzheimer disease. Front Neurol 4:122.

Collins GA and Goldberg AL (2017) The Logic of the 26S Proteasome. Cell 169:792–806.

Cuervo AM, Stefanis L, Fredenburg R, Lansbury PT and Sulzer D (2004) Impaired degradation of mutant alpha-synuclein by chaperone-mediated autophagy. Science 305:1292–1295.

D’Arcy P, Brnjic S, Olofsson MH, Fryknas M, Lindsten K, De Cesare M, Perego P, Sadeghi B, Hassan M, Larsson R and Linder S (2011) Inhibition of proteasome deubiquitinating activity as a new cancer therapy. Nat Med 17:1636–1640.

de Poot SAH, Tian G and Finley D (2017) Meddling with Fate: The Proteasomal Deubiquitinating Enzymes. J Mol Biol.

Finley D, Chen X and Walters KJ (2016) Gates, Channels, and Switches: Elements of the Proteasome Machine. Trends Biochem Sci 41:77–93.

Glickman MH and Ciechanover A (2002) The ubiquitin-proteasome proteolytic pathway: destruction for the sake of construction. Physiol Rev 82:373–428.

Goitea VE and Hallak ME (2015) Calreticulin and Arginylated Calreticulin Have Different Susceptibilities to Proteasomal Degradation. J Biol Chem 290:16403–16414.

Hanna J, Hathaway NA, Tone Y, Crosas B, Elsasser S, Kirkpatrick DS, Leggett DS, Gygi SP, King RW and Finley D (2006) Deubiquitinating enzyme Ubp6 functions noncatalytically to delay proteasomal degradation. Cell 127:99–111.

Hassiepen U, Eidhoff U, Meder G, Bulber JF, Hein A, Bodendorf U, Lorthiois E and Martoglio B (2007) A sensitive fluorescence intensity assay for deubiquitinating proteases using ubiquitin-rhodamine110-glycine as substrate. Anal Biochem 371:201–207.

Homan CC, Kumar R, Nguyen LS, Haan E, Raymond FL, Abidi F, Raynaud M, Schwartz CE, Wood SA, Gecz J and Jolly LA (2014) Mutations in USP9X are associated with X-linked intellectual disability and disrupt neuronal cell migration and growth. Am J Hum Genet 94:470–478.

Homma T, Ishibashi D, Nakagaki T, Fuse T, Mori T, Satoh K, Atarashi R and Nishida N (2015) Ubiquitin-specific protease 14 modulates degradation of cellular prion protein. Sci Rep 5:11028.

Hyrskyluoto A, Bruelle C, Lundh SH, Do HT, Kivinen J, Rappou E, Reijonen S, Waltimo T, Petersen A, Lindholm D and Korhonen L (2014) Ubiquitin-specific protease-14 reduces cellular aggregates and protects against mutant huntingtin-induced cell degeneration: involvement of the proteasome and ER stress-activated kinase IRE1alpha. Hum Mol Genet 23:5928–5939.

Jacobson AD, MacFadden A, Wu Z, Peng J and Liu CW (2014) Autoregulation of the 26S proteasome by in situ ubiquitination. Mol Biol Cell 25:1824–1835.

Jacobson AD, Zhang NY, Xu P, Han KJ, Noone S, Peng J and Liu CW (2009) The lysine 48 and lysine 63 ubiquitin conjugates are processed differently by the 26 s proteasome. J Biol Chem 284:35485–35494.

Jin YN, Chen PC, Watson JA, Walters BJ, Phillips SE, Green K, Schmidt R, Wilson JA, Johnson GV, Roberson ED, Dobrunz LE and Wilson SM (2012) Usp14 deficiency increases tau phosphorylation without altering tau degradation or causing tau-dependent deficits. PLoS One 7:e47884.

Kim HT and Goldberg AL (2017) The deubiquitinating enzyme Usp14 allosterically inhibits multiple proteasomal activities and ubiquitin-independent proteolysis. J Biol Chem 292:9830–9839.

Kiprowska MJ, Stepanova A, Todaro DR, Galkin A, Haas A, Wilson SM and Figueiredo-Pereira ME (2017) Neurotoxic mechanisms by which the USP14 inhibitor IU1 depletes ubiquitinated proteins and Tau in rat cerebral cortical neurons: Relevance to Alzheimer’s disease. Biochimica et biophysica acta 1863:1157–1170.

Kish-Trier E and Hill CP (2013) Structural biology of the proteasome. Annu Rev Biophys 42:29–49.

Komander D and Rape M (2012) The ubiquitin code. Annual review of biochemistry 81:203–229.

Koulich E, Li X and DeMartino GN (2008) Relative structural and functional roles of multiple deubiquitylating proteins associated with mammalian 26S proteasome. Mol Biol Cell 19:1072–1082.

Lam YA, Xu W, DeMartino GN and Cohen RE (1997) Editing of ubiquitin conjugates by an isopeptidase in the 26S proteasome. Nature 385:737–740.

Lee BH, Finley D and King RW (2012) A High-Throughput Screening Method for Identification of Inhibitors of the Deubiquitinating Enzyme USP14. Curr Protoc Chem Biol 4:311–330.

Lee BH, Lee MJ, Park S, Oh DC, Elsasser S, Chen PC, Gartner C, Dimova N, Hanna J, Gygi SP, Wilson SM, King RW and Finley D (2010) Enhancement of proteasome activity by a small-molecule inhibitor of USP14. Nature 467:179–184.

Lee MJ, Lee BH, Hanna J, King RW and Finley D (2011) Trimming of ubiquitin chains by proteasome-associated deubiquitinating enzymes. Mol Cell Proteomics 10:R110 003871.

Leggett DS, Hanna J, Borodovsky A, Crosas B, Schmidt M, Baker RT, Walz T, Ploegh H and Finley D (2002) Multiple associated proteins regulate proteasome structure and function. Mol Cell 10:495–507.

Li J, Wang J, Hou W, Jing Z, Tian C, Han Y, Liao J, Dong MQ and Xu X (2011) Phosphorylation of Ataxin-10 by polo-like kinase 1 is required for cytokinesis. Cell Cycle 10:2946–2958.

Liu B, Jiang S, Li M, Xiong X, Zhu M, Li D, Zhao L, Qian L, Zhai L, Li J, Lu H, Sun S, Lin J, Lu Y, Li X and Tan M (2018) Proteome-wide analysis of USP14 substrates revealed its role in hepatosteatosis via stabilization of FASN. Nat Commun 9:4770.

Ortuno D, Carlisle HJ and Miller S (2016) Does inactivation of USP14 enhance degradation of proteasomal substrates that are associated with neurodegenerative diseases? F1000Res 5:137.

Peth A, Besche HC and Goldberg AL (2009) Ubiquitinated proteins activate the proteasome by binding to Usp14/Ubp6, which causes 20S gate opening. Mol Cell 36:794–804.

Peth A, Kukushkin N, Bosse M and Goldberg AL (2013) Ubiquitinated proteins activate the proteasomal ATPases by binding to Usp14 or Uch37 homologs. J Biol Chem 288:7781–7790.

Randles L, Anchoori RK, Roden RB and Walters KJ (2016) The Proteasome Ubiquitin Receptor hRpn13 and Its Interacting Deubiquitinating Enzyme Uch37 Are Required for Proper Cell Cycle Progression. J Biol Chem 291:8773–8783.

Saeki Y (2017) Ubiquitin recognition by the proteasome. J Biochem 161:113–124.

Sell GL and Margolis SS (2015) From UBE3A to Angelman syndrome: a substrate perspective. Front Neurosci 9:322.

Smibert P, Yang JS, Azzam G, Liu JL and Lai EC (2013) Homeostatic control of Argonaute stability by microRNA availability. Nat Struct Mol Biol 20:789–795.

Sreedharan J, Blair IP, Tripathi VB, Hu X, Vance C, Rogelj B, Ackerley S, Durnall JC, Williams KL, Buratti E, Baralle F, de Belleroche J, Mitchell JD, Leigh PN, Al-Chalabi A, Miller CC, Nicholson G and Shaw CE (2008) TDP-43 mutations in familial and sporadic amyotrophic lateral sclerosis. Science 319:1668–1672.

Stamos JL and Weis WI (2013) The beta-catenin destruction complex. Cold Spring Harb Perspect Biol 5:a007898.

Tomko RJ, Jr. and Hochstrasser M (2013) Molecular architecture and assembly of the eukaryotic proteasome. Annual review of biochemistry 82:415–445.

Verma R, Aravind L, Oania R, McDonald WH, Yates JR, 3rd, Koonin EV and Deshaies RJ (2002) Role of Rpn11 metalloprotease in deubiquitination and degradation by the 26S proteasome. Science 298:611–615.

Wilkinson KD, Urban MK and Haas AL (1980) Ubiquitin is the ATP-dependent proteolysis factor I of rabbit reticulocytes. J Biol Chem 255:7529–7532.

Wobst HJ, Wesolowski SS, Chadchankar J, Delsing L, Jacobsen S, Mukherjee J, Deeb TZ, Dunlop J, Brandon NJ and Moss SJ (2017) Cytoplasmic Relocalization of TAR DNA-Binding Protein 43 Is Not Sufficient to Reproduce Cellular Pathologies Associated with ALS In vitro. Front Mol Neurosci 10:46.

Yao T and Cohen RE (2002) A cryptic protease couples deubiquitination and degradation by the proteasome. Nature 419:403–407.

Yao T, Song L, Xu W, DeMartino GN, Florens L, Swanson SK, Washburn MP, Conaway RC, Conaway JW and Cohen RE (2006) Proteasome recruitment and activation of the Uch37 deubiquitinating enzyme by Adrm1. Nat Cell Biol 8:994–1002.

